# MissenseHMM: state-based annotations for missense variants through joint modeling of pathogenicity scores

**DOI:** 10.64898/2026.01.31.703062

**Authors:** Runjia Li, Jason Ernst

**Author notes:** Correspondence (JE).

## Abstract

Many computational predictors of missense variant pathogenicity are available. To capture information across various predictors, we propose MissenseHMM, which learns states corresponding to combinatorial patterns of variant prioritizations. We applied MissenseHMM to 43 predictors, annotating over 70 million missense variants with 20 states that showed distinct predictor scores patterns, amino acid substitutions and other genomic annotation enrichments. MissenseHMM state annotations enhanced individual predictors’ associations with clinical pathogenic variants and deep mutational scanning data, and also provided insight into the performances of various protein language models. Overall, MissenseHMM complements pathogenicity predictors and is an annotation resource for missense variant interpretation.

## Background

Interpreting missense variants, which cause single-mutation changes to the amino acid sequences of proteins, is a significant challenge. Despite ongoing efforts to document their clinical implications^1,2^, a large portion of missense variants have uncertain phenotypic impact. With whole exome sequencing (WES) and whole genome sequencing (WGS) leading to large-scale identification of missense variants^2–4^, the ability to effectively annotate and characterize missense variants is a limiting factor in many genetic studies attempting to link missense variants to disease^5^.

To address this, many computational predictors have been proposed to predict the disease impact of missense variants^6,7^. These predictors leverage information about missense variants such as their local DNA sequence, conservation, or epigenomic and related annotations to generate missense pathogenicity scores of predicted deleteriousness. A number of predictors are based on integrating various types of features in supervised training frameworks^8–13^. Recent advances in deep learning have led to the emergence of protein language models^14–17^ that directly infer the impact of missense variants based on only sequence information.

While various missense scores are available, there remain substantial differences among scores. For instance in the dbNSFP v 4.0 database^6,7^, which provides scores from more than 40 pathogenicity predictors for a comprehensive collection of missense variants in the human genome, about 56% of the pairwise correlations between scores are smaller than 0.5^7^. These differences could be due to the scores’ differences in input features or methods of generating predictions from the features. Recognizing differences among different scores, ensemble-based methods have been developed that integrate scores from multiple predictors into a single predictor for improving pathogenicity prediction^8–11,18–25^. However, such ensemble approaches still produce univariate scores that convey limited information about what was captured by the different scores.

To provide an integrative annotation of missense variants that provides additional interpretable information summarizing information in the individual missense scores, we develop MissenseHMM. MissenseHMM uses a multivariate hidden Markov model (HMM) to infer missense variant prioritization states, which correspond to combinatorial patterns of the variant prioritization of different scores while considering the spatial context. This is analogous to approaches that have been used to annotate genomes based on combinatorial and spatial patterns of multiple epigenetic marks^26,27^. For each possible missense variant, MissenseHMM annotates it to one state. We implement MissenseHMM as an extension of the ChromHMM^26^ software. We apply MissenseHMM based on 43 scores curated by the dbNSFP database to annotate more than 70 million possible human missense variants. We show that the different MissenseHMM states are associated with distinct properties of missense scores and other genomic features. The state annotations also provide additional information complementary to individual scores that can improve their prediction of the variants’ pathogenicity status. We expect MissenseHMM annotations to be a useful resource for analyzing missense variants.

## Results

### MissenseHMM for discovering and annotating missense variant prioritization states

We developed MissenseHMM, an approach to annotate each missense variant using states learned from a multivariate Hidden Markov Model (HMM) based on the combinatorial patterns of variants prioritized by various missense pathogenicity scores and their spatial context (Figure 1). MissenseHMM first binarizes the scores for each missense variant. For each score, the top *Q*% missense variants are encoded as present and the remaining scored variants are encoded as absent, where *Q* is a user-specified parameter (Methods). MissenseHMM then generates one training sequence for each gene by randomly selecting one missense variant for each unique genomic position within the coding portion of its exons where a missense variant can occur, and then concatenating them. From these training sequences and a specified number of states, MissenseHMM learns in an unsupervised manner the HMM parameters (Methods).

**Figure 1:**
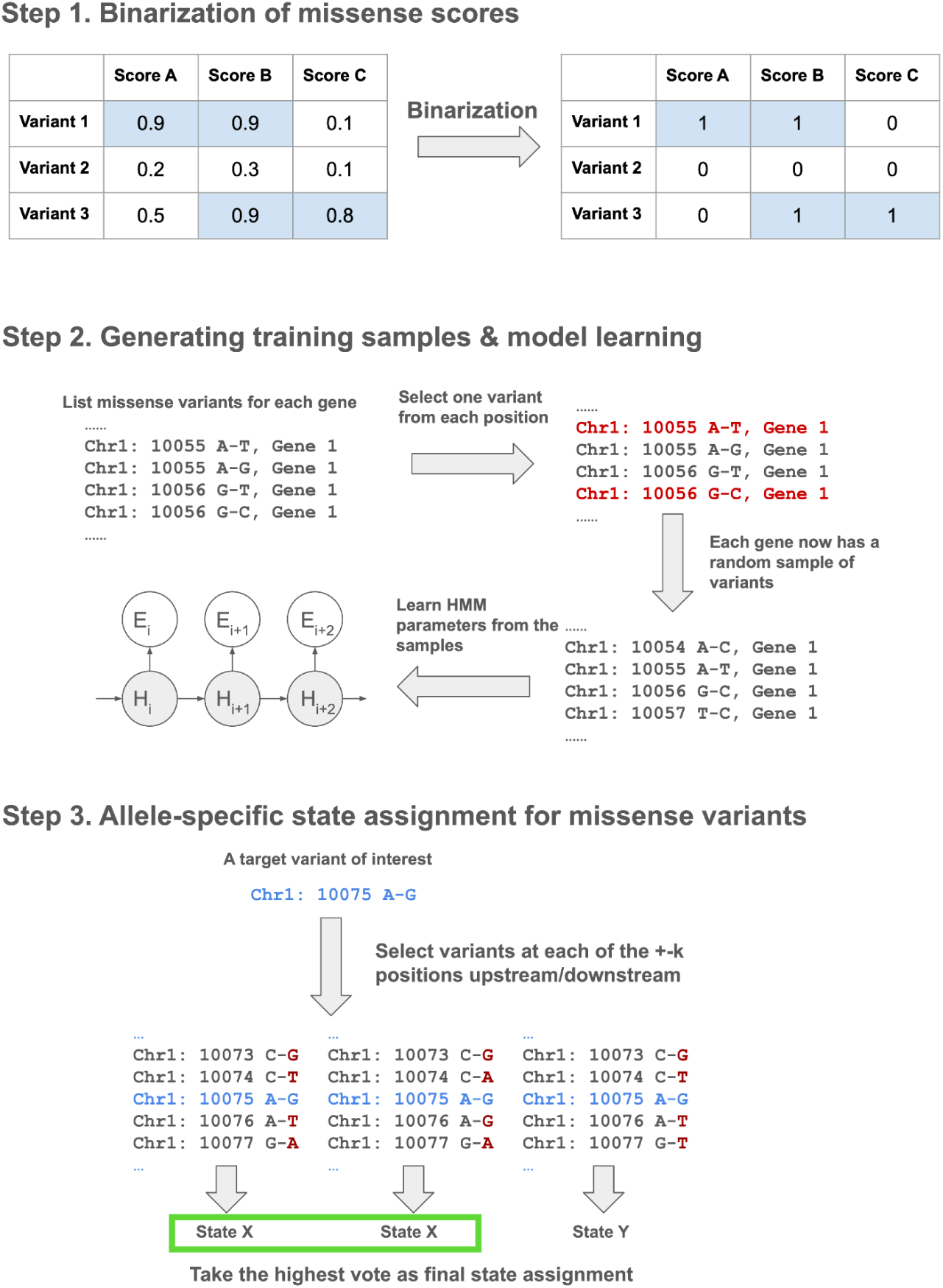
Schematic of the MissenseHMM model training and state assignment pipeline. Step 1: For each missense variant, MissenseHMM binarizes each missense score based on whether it is in the top *Q*% across all missense variants. Step 2: For each gene, MissenseHMM generates a random sample by selecting a random missense variant at each unique position for which there is at least one missense variant. MissenseHMM then trains an HMM where there is a sample from each gene. Step 3: For a target missense variant of interest, MissenseHMM generates *N* combinations of missense variants *k* positions upstream and downstream. For each combination, MissenseHMM gives the target variant an HMM state assignment. MissenseHMM then takes as the final state assignment for the target variant a state assignment occurring most frequently among the combinations.

After a model is learned, to efficiently assign a state to each possible variant, MissenseHMM employs a local window-based approach^24,25^ (Methods). Specifically, for each target variant, MissenseHMM generates *N* sequences by randomly selecting missense variants for each of the *k* positions upstream and downstream of the target variant. MissenseHMM then for each sequence assigns a state to the target variant and retains a mode assignment across sequences as the final assignment for the variant (Methods).

### MissenseHMM learns distinct states from scores curated in dbNSFP

We applied MissenseHMM using 43 missense variant prioritization scores (Table S1) to annotate 70,842,617 missense variants in the dbNSFP database v5.0a^6,7^ for the human hg38 assembly. We trained MissenseHMM models using 10 to 50 states in increments of 10 and set *Q*=10, *k*=3 and *N*=9. Considering a trade-off between ease of state interpretation and additional information gained (Methods), we focused our subsequent analysis on the 20-state model. We first broadly categorized the states based on how many scores its corresponding emission probability was relatively high in the state. To do so, we first for each score ranked the states in descending order based on the emission probability for that score. Then for each state we counted the number of scores in which its emission probability ranked among the top five across all states (subsequently referred to as “top-5 scores”). Based on these counts, we ordered the states and then categorized them into four groups (Figure 2a, Table S2). Group 1 comprises three states: 1, 2 and 3, representing missense variants prioritized by the most missense scores, with each state having at least 35 top-5 scores. Notable distinctions between these states include State 3 having lower emissions for the four VARITY^30^-based scores, which were the only scores trained on deep mutational scanning (DMS) data and that State 2 has lower emissions for DEOGEN2^31^ and four ensemble-based scores (MVP^11^, MetaSVM^9^, MetaLR^24^ and M-CAP^10^). Group 2 comprises states 4-9, which had relatively high emission parameter values for an intermediate number of scores, each having between 7 and 18 top-5 scores. For example, State 7 has the highest emission for PrimateAI^13^ and MPC^12^ across all states and also exhibits relatively high emissions for two conservation scores (Table S1): PhastCons17way_primate^32^ (2nd highest across all states) and LIST-S2^33^ (4th), and several other scores that use sequencing alignment-based features, including gMVP^34^ (4th), PHACTboost^30^ (4th), and AlphaMissense^15^ (4th). Group 3 comprises states 10-18, each with between 1-5 top-5 scores, in other words showing specificity to a limited number of scores. For example, State 18 has the highest emission for MutFormer^16^ across all states but has low emissions for all other scores, with an average emission rank of 12.7. Group 4 comprises State 19 and State 20, both of which have low emissions across all scores and no top-5 scores. More than 40% of the missense variants are assigned to these two states. Overall, the states exhibit a wide variety of patterns of variant prioritization across the scores.

**Figure 2:**
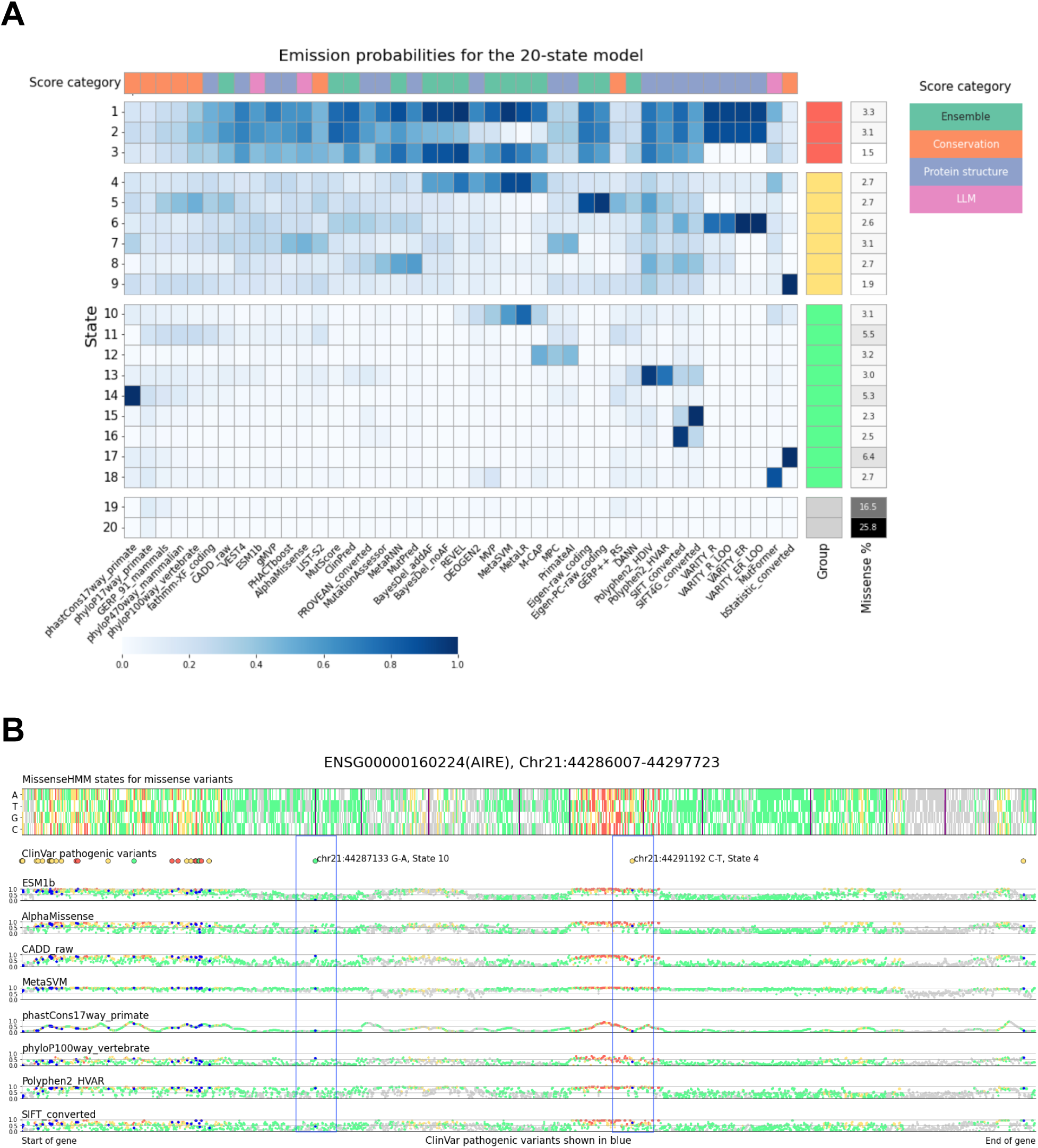
Overview of the 20-state MissenseHMM model. **(A)** Emission probabilities of the 20-state MissenseHMM model. From top to bottom the states are ordered and numbered based on the number of scores in which their emission probability ranked among the top five across all states. The right color bar indicates the state grouping and the percentage of missense variants assigned to each state. The top color bar indicates the broad categorizations of the scores. The individual emission values corresponding to the table are in Table S2. **(B)** An example of a gene (AIRE) annotated using MissenseHMM states. The exons of the gene are concatenated. The top heatmap shows the state assignments for each missense variant, using the same color scheme as the groupings in (A). Synonymous variants are shown as white. Vertical lines separate exons. The locations of the ClinVar pathogenic variants are shown below, also with the same color scheme indicating the state assignments for these variants. The eight scatterplots below show selected individual predictor score values for the missense variants, scaled between 0 and 1 (Methods). The same color scheme is used except for ClinVar pathogenic variants which are now for clarity colored blue. In this example, two ClinVar pathogenic variants show different prioritization patterns across the selected scores. Chr21: 44287133 G-A has low scores for the two LLM-based predictors (bottom 9% for ESM1b and bottom 22% for AlphaMissense) but high scores for others, such as MetaSVM (0.93), for which its state annotation, State 10, had high emission parameter values. Chr21: 44291192 C-T has low conservation scores (bottom 24% for phastCons17way_primate and bottom 33% for phyloP100way_vertebrate) but high scores for others, reflecting that its assigned state, State 4, has the lowest average emissions for conservation scores (0.16) among the four categories of scores.

To confirm that MissenseHMM’s consideration of spatial context through the use of transition parameters (Table S3) has an effect on variant annotations, we compared MissenseHMM’s 20-state model to an equivalent model but with uniform transition probabilities between states.

On average, each variant in the original 20-state model had a 22% chance of being assigned to a different state in the uniform transition model (ranging from 3% to 69% across the different states, Figure S2). Additionally, the original model produced longer segments of consecutive missense variants assigned to the same state, compared to the uniform transition model (5.7 million segments, with an average length of 4.8 variants, compared to 6.0 million segments with an average length of 3.8 variants; one-sided Mann-Whitney U p-value < 10^-300^). These results suggest that considering the sequential properties between the variants can capture additional information.

### MissenseHMM states show distinct enrichment for external annotations

To better characterize the biological properties of the MissenseHMM states, we computed enrichments of their corresponding positions with several external genomic annotations (Methods). We first computed enrichments of the missense variants assigned to a state for CpG islands, transcription start sites (TSS), transcription end sites (TES), and 2kb windows surrounding TSS, using all missense variants as background (Figure 3A). The highest enrichment for any combination of external annotation and state (9.2 fold) was between TSS and State 8, the state with the third-highest emission for MutPred^21^. State 16, the state with the highest emission for SIFT, also had a notably high enrichment for TSS (4.9 fold). A different state, State 12, had the highest enrichment for CpG islands and the 2kb windows surrounding TSS (6.0 and 3.5 fold, respectively), while not being enriched at TSS (0.8 fold). State 12 had relatively low emissions for most scores while having the second-highest emission for MPC and PrimateAI and the fourth-highest emission for M-CAP. The three states with the highest enrichment for TES (1.6 fold) were States 9 and 17, which had the second-highest and highest emission probability for bStatistic, respectively, and State 16, which had the highest emission probability for SIFT.

**Figure 3:**
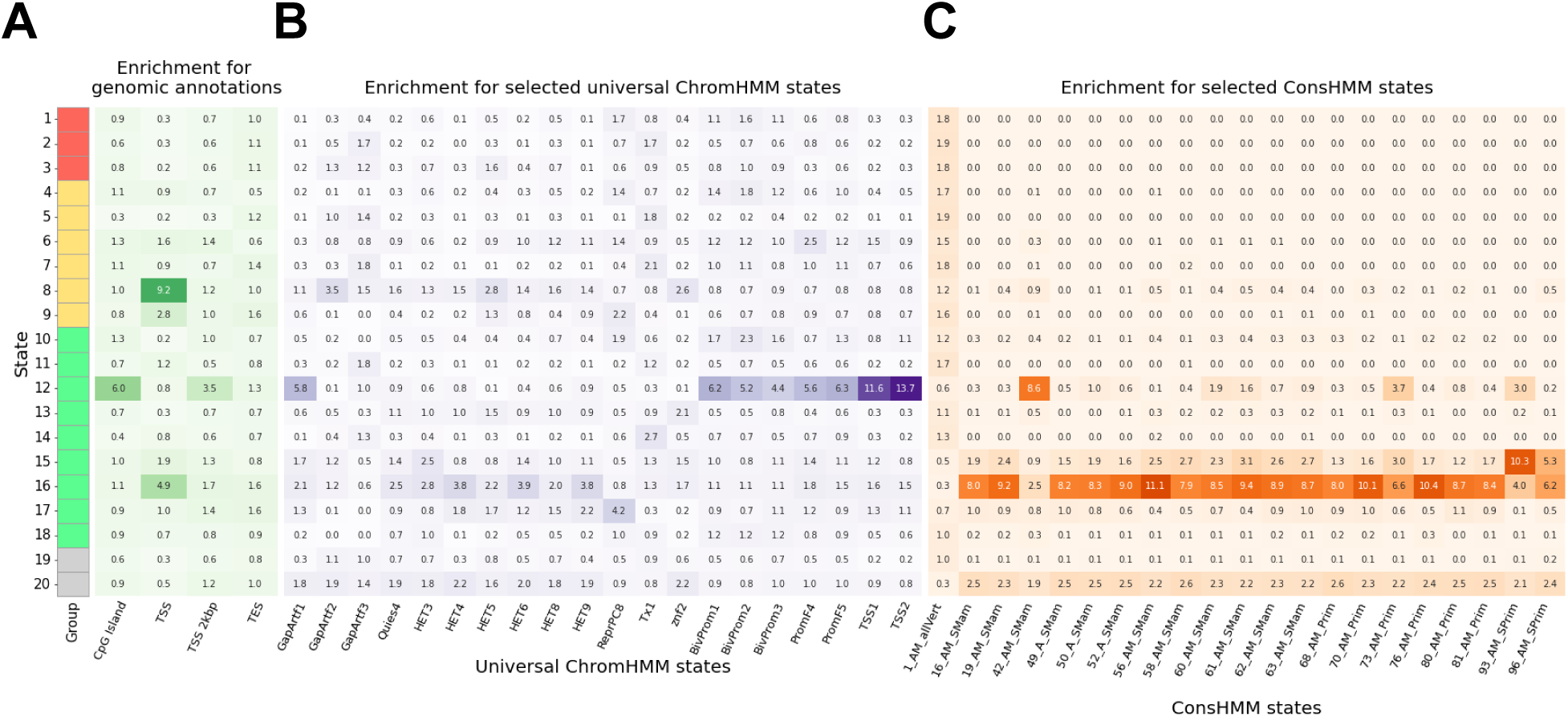
MissenseHMM states’ overlap enrichments for external annotations. **(A)**: MissenseHMM state enrichments for genomic annotations including CpG islands, TSS, TES, and 2kb windows surrounding TSS. **(B, C)**: Enrichments for the top-20 universal ChromHMM **(B)** and ConsHMM states **(C)** with the highest variances of enrichments across the MissenseHMM states. For ConsHMM, the state with the highest extent of alignment and matching across all vertebrate species (1_AM_allVert) is also shown. The colored bar on the left indicates the state group. For universal ChromHMM states the mnemonics as defined by Vu and Ernst^36^ are GapArtf: assembly gaps and artifacts. Quies: quiescent; HET: heterochromatin; ReprPC: polycomb repressed; Tx: transcribed; Znf: zinc finger genes; BivProm: bivalent promoter; PromF: flanking promoter; TSS: transcription start site. For ConsHMM states the mnemonics are defined following Arneson and Ernst^29^. AM_allVert: the state has high align and match probabilities for all vertebrates. AM_SMam: the state has high align and match probabilities for mammals, but missing notable subsets. A_SMam: the state has high align probabilities for mammals, but low match probabilities. AM_Prim: the state has high align and match probabilities for primates. AM_SPrim: the state has high align and match probabilities for a few subsets of primates. The enrichment for all universal ChromHMM states and ConsHMM can be found in Table S4 and Table S5, respectively.

Next, we analyzed the enrichment of MissenseHMM states for annotations from the 100-state full-stack universal ChromHMM state model^32^. These ChromHMM state annotations represent combinatorial and spatial patterns of chromatin marks across more than 1000 data sets from over 100 cell and tissue types (Figure 3B, Table S4) and capture diverse classes of genomic elements. State 12, the state most enriched for CpG islands and regions within 2kb of a TSS, also showed the strongest enrichment for any chromatin state, 13.7 fold for the TSS2, using the same missense background as above. Additionally it showed strong enrichments for TSS1 (11.6 fold), which along with TSS2 are associated with promoter regions in addition to TSS, along with other promoter associated chromatin states (4.4-6.2 fold enriched for three bivalent promoter states (BivProm1, 2, 3); 5.6-6.3 fold enriched for two flanking promoter states (PromF4, 5)) consistent with its enrichment for 2kb windows surrounding TSS. State 12 also had a 5.8-fold enrichment for a chromatin state associated with assembly gaps or alignment artifacts (GapArtf1). The state that showed the next highest enrichment (4.2 fold) for any chromatin state was State 17, the state with the highest emission for bStatistic, for one of the polycomb repressed states, ReprPC8.

We also computed enrichments of MissenseHMM states for annotations from a 100-state ConsHMM model^28,29^ (Figure 3C, Table S5). The ConsHMM state annotations from this model annotate patterns of sequence conservation among 100 vertebrate species in terms of nucleotide alignment and matching the human reference genome. Among the MissenseHMM states, five states (12, 15, 16, 17 and 20) depleted for ConsHMM State 1, the state with the highest level of alignment and matching throughout the vertebrates. These five states also showed enrichments for many of the other of the 100 ConsHMM states associated with less constraint, 41 to 99 states in total, in contrast the other 15 MissenseHMM states enriched for between 0 and 5 other states. Among the five states, State 12 had the highest enrichment for ConsHMM State 42 (8.6 fold), a state with moderate alignment and matching signal across many vertebrates, and also previously shown to be highly enriched for CpG islands and TSS^29^. This state was also the state with the lowest average emissions for conservation scores.

### MissenseHMM states show distinct preferences for amino acid substitutions

We next investigated if the MissenseHMM states show different enrichments for specific amino acid substitutions. To do this, we counted the number of each amino acid substitution for missense variants assigned to each state and compared it with the total counts across all states. We observed that states in Groups 1 and 2 tended to have a greater relative frequency than average for the more impactful substitutions as predicted based on BLOSUM62^37^ values, whereas states in Groups 3 and 4 tended to show a lower relative frequency of those substitutions (Figure 4). Within Group 1, States 1 and 2 showed similar preferences across all substitutions (Spearman correlation 0.93), differing from that of State 3 (Spearman correlation 0.48 and 0.42, respectively). Meanwhile, State 6 from Group 2 exhibited a highly similar pattern as States 1 and 2 (Spearman correlation 0.85 and 0.87, respectively). Notably, States 1, 2 and 6 all had higher emissions for the four VARITY-based scores, (average emission 0.88-0.92) compared to State 3 (average emission 0.00003). Despite the overall lower relative frequency for impactful substitutions among states in Group 3, some states in Group 3 did show a high relative frequency for specific types of impactful substitutions. An example of this being States 12, 13, and 16 for proline to leucine (P-L), for which their relative frequency ranked 3rd-5th among all states (with 0.5-0.7% greater relative frequency, corresponding to a 1.4 to 1.7 fold enrichment), suggesting that the score-specific states still associate with impactful substitutions predicted based on BLOSUM62 values.

**Figure 4:**
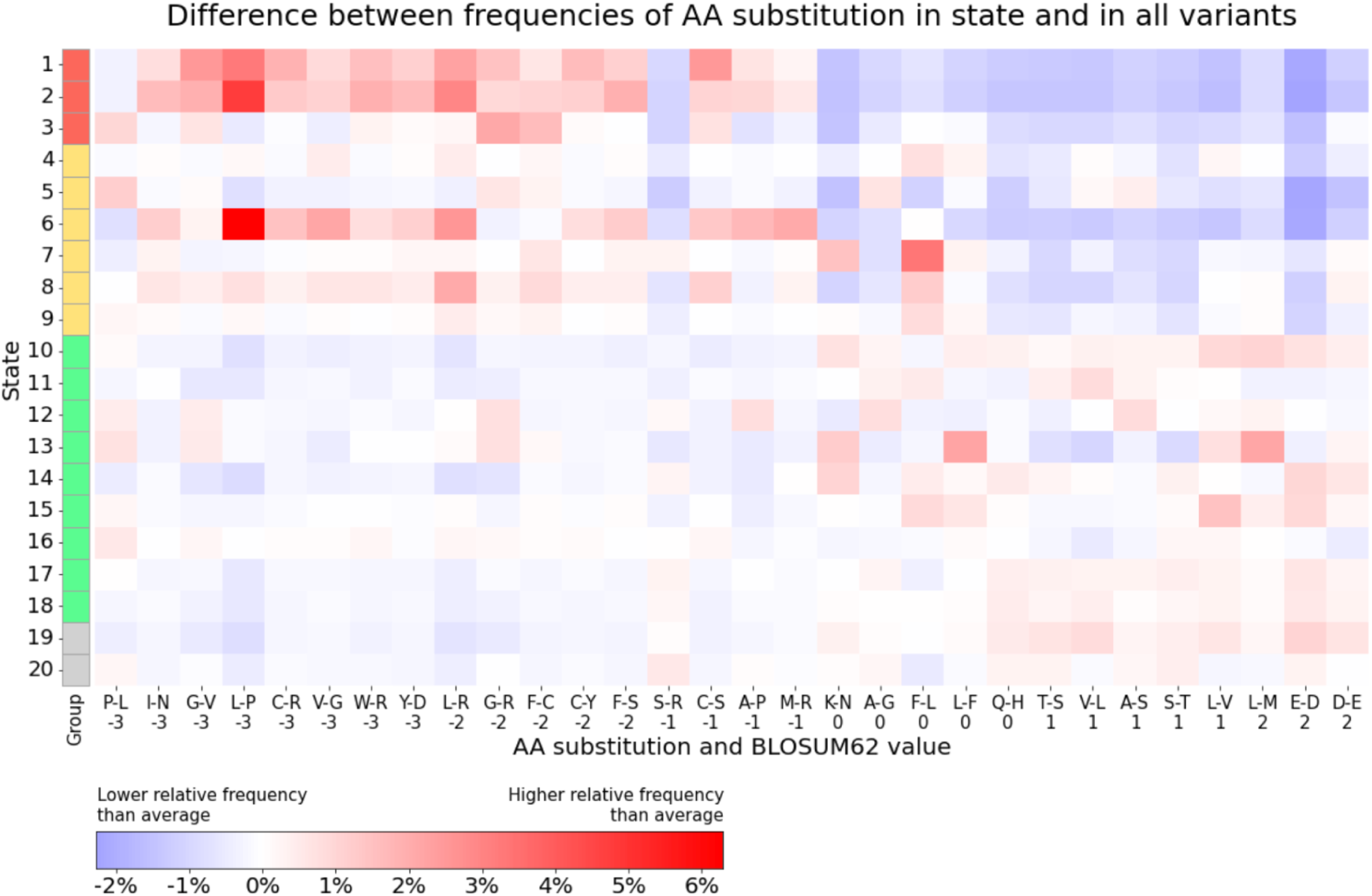
MissenseHMM states’ preference for selected amino acid substitutions. Each row represents a state, and each column corresponds to a specific amino acid (AA) substitution. The rows are ordered based on state groupings and the columns based on the substitutions’ BLOSUM62 value. Positive values indicate more likely, and thus less impactful, substitutions, while negative values indicate the opposite. The value in each cell denotes the difference between the proportion of that substitution in the given state and its total proportion across all states. Positive values are colored red, and negative values are colored blue. The top 30 amino acid substitutions with the largest variances across states are shown.

### MissenseHMM annotations contain complementary information predictive of pathogenic variants

We next investigated if MissenseHMM annotations contain complementary information to individual missense scores predictive of pathogenic variants. For this, we collected annotated pathogenic variants from the ClinVar^1^ database as the positive set and singleton (MAC = 1) variants from the gnomAD^2^ database as the negative set, mirroring evaluation sets used previously in the analysis of pathogenic variant prediction accompanying release of dbNSFP v4.0^7^. We note that a caveat of the analysis is that some of the scores in the MissenseHMM model already used ClinVar or gnomAD variants in their training (Table S6), which prevents a fully unbiased evaluation of predictiveness for previously unseen variants.

We observed that even for variants highly ranked by a missense score the probability of a variant being labelled pathogenic in ClinVar can vary substantially depending on the variant’s MissenseHMM state. For instance, a variant with a scaled ESM1b score greater than 0.90, corresponding to the top 10% of the raw ESM1b scores, would have a 95.4% probability of being a ClinVar-labelled pathogenic variant if it were in State 1, a 77.1% probability of being positive in State 18, but only a 39.4% probability of being positive in State 8 (Figure 5, Table S7). Overall this probability has a mean of 41% and a standard deviation of 31% across the states. These findings suggest that MissenseHMM states can provide additional information about the pathogenicity of missense variants beyond what is captured by individual scores.

**Figure 5:**
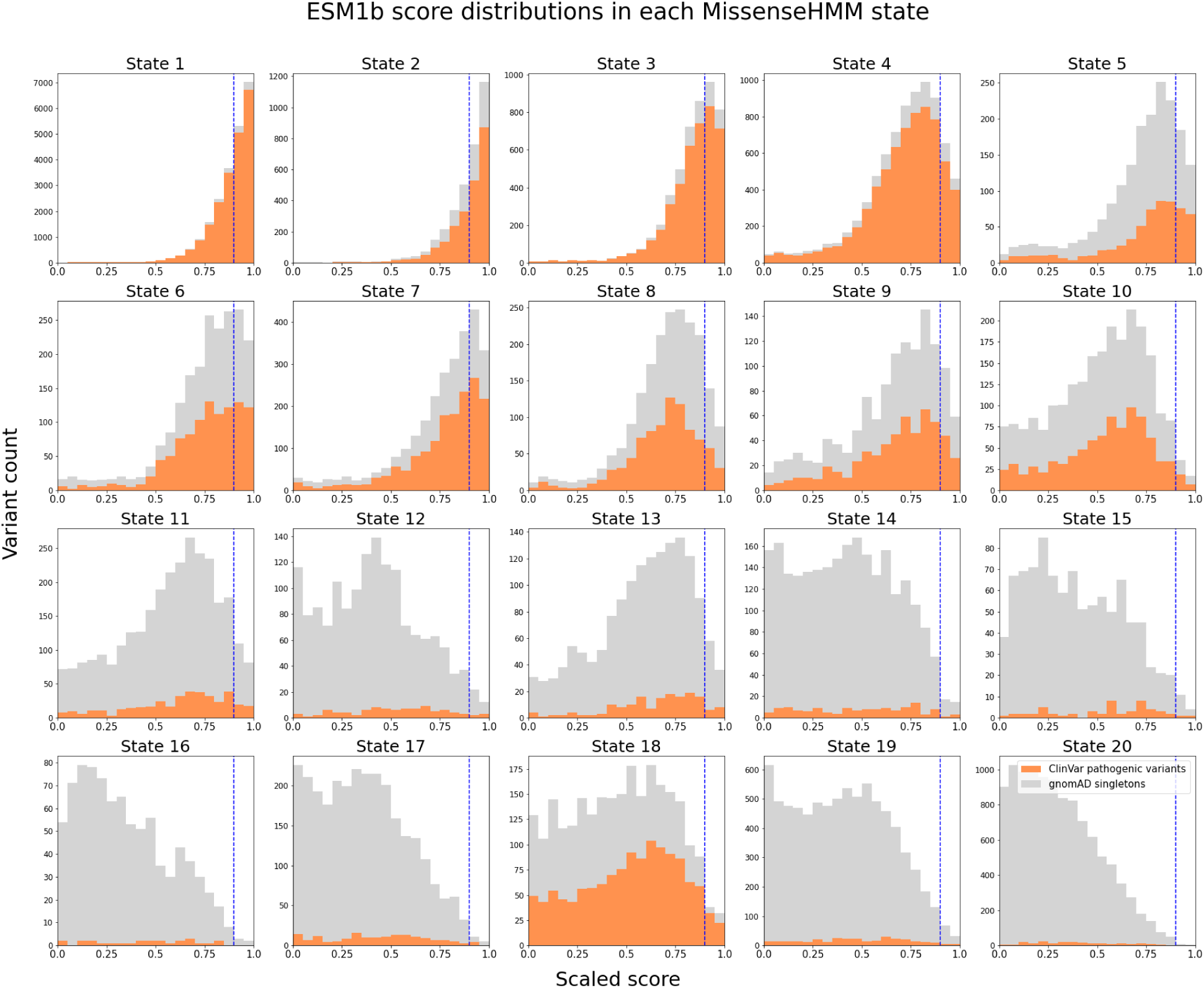
ESM1b score distributions for ClinVar pathogenic variants and gnomAD singletons in each MissenseHMM state. The MissenseHMM states are ordered by their groups. The ESM1b score on the x-axis has been scaled between 0 and 1, with 1 indicating highest disease impact. Histogram bars for gnomAD singletons (grey) are stacked on top of those for ClinVar pathogenic variants (orange). Y-axis shows the number of variants in each histogram bar, with different scales in each sub-histogram. Vertical dashed lines represent scaled score = 0.90. Full results for all feature scores of MissenseHMM can be found in Table S7.

We next systematically evaluated the extent to which combining MissenseHMM annotations with individual scores could potentially lead to increased pathogenicity predictions compared to using individual scores alone. For this, we first quantified the performance of each of the 43 missense scores in distinguishing ClinVar pathogenic variants from gnomAD singletons alone using a logistic regression model with the score as its sole feature. We then compared its performance with a logistic regression model that also had binary features corresponding to the presence of each MissenseHMM state. The logistic regression models were trained to discriminate ClinVar pathogenic variants from gnomAD singletons (Methods). On average, the incorporation of the MissenseHMM states led to a 16% improvement in AUROC, ranging from 57% (for phyloP 17-way primate) to 2% (PHACTboost) (Figure 6). The percent improvement was largest for conservation scores (37% on average) and smallest for ensemble scores (8.7% on average). Overall, these results suggest MissenseHMM annotations may provide information to increase the predictive power of individual missense scores of curated pathogenic variants.

**Figure 6:**
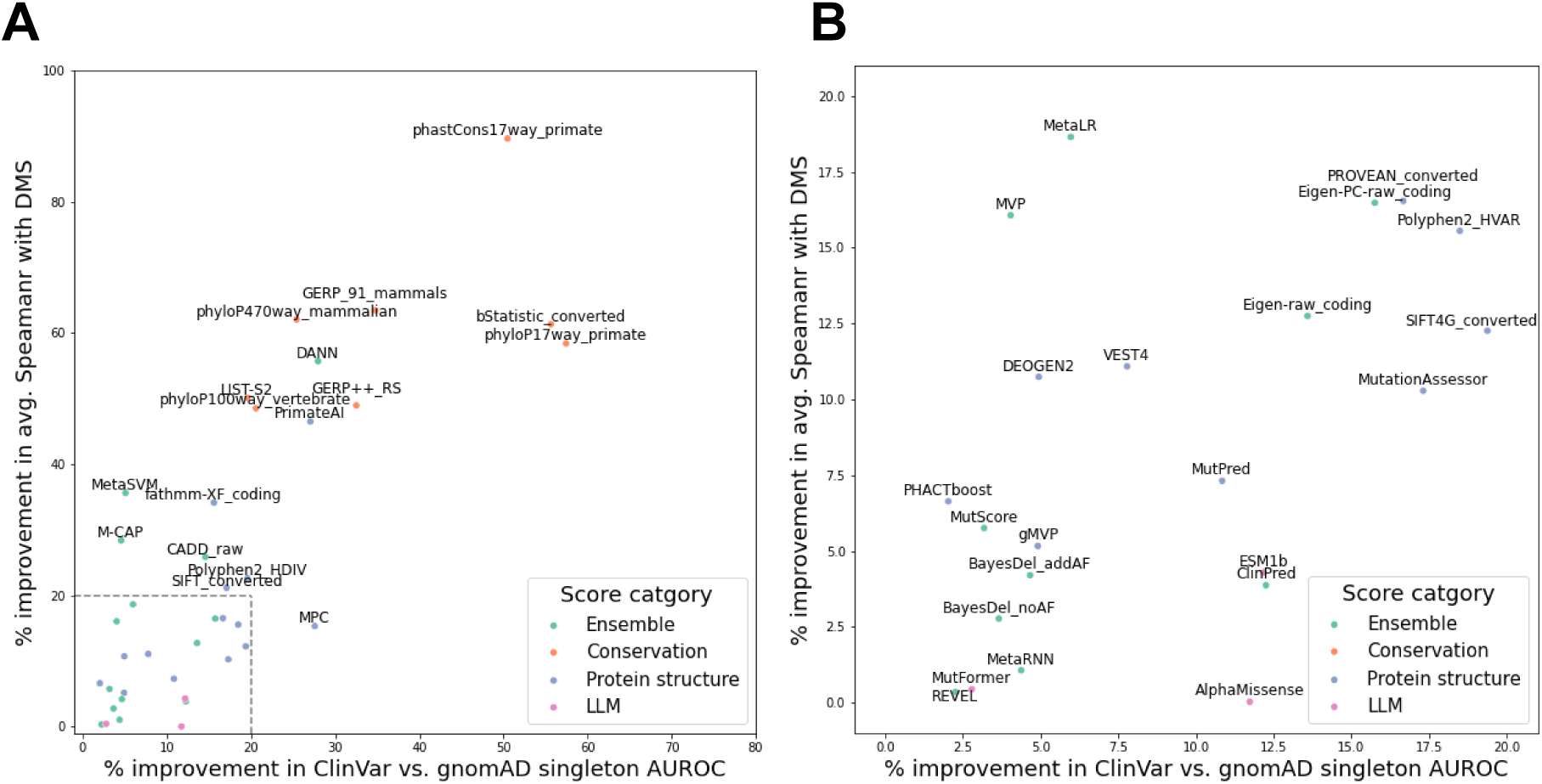
Additional predictive information through integration of MissenseHMM state annotations with missense scores of ClinVar pathogenic variants and of DMS experimental measurements, with and without integration of MissenseHMM states. The x-axis shows % improvement in AUROC using a logistic regression model to predict ClinVar pathogenic variants vs. gnomAD singletons, from using each score as a feature to using each score and the MissenseHMM state as features. The y-axis shows % improvement in average Spearman correlation using a linear regression model to predict DMS experimental measurements, from using each score as a feature to using each score and the MissenseHMM state as features. The scores are colored based on their broad category. **(A)** The full set of scores is labelled, except for the scores in the box in the lower-left corner. **(B)** Zoomed-in view of the lower-left corner in **A**.

### MissenseHMM annotations contain complementary information predictive of DMS data

We additionally evaluated the extent to which MissenseHMM annotations contain complementary information to individual scores that are predictive of experimentally measured functional effects of variants. To do so, we utilize DMS experiments, which are based on assays that measure the functional impact of specific mutations on proteins. Specifically, we obtained a set of precompiled DMS experimental data for 36 proteins from Livesey and Marsh^38^, with the experimental data for each protein potentially containing more than one measurement. Since VARITY scores were directly trained on DMS datasets, we excluded the 10 proteins used in their training data. This resulted in 26 proteins with 64 measurements. For each measurement, we compared the predictive performance of each missense score, excluding the four VARITY-based scores, using linear regression models with and without the MissenseHMM state assignment as an additional categorical feature (Methods). We observed that all remaining missense scores showed improvement in Spearman correlation between their predicted and true DMS measurements after the integration of MissenseHMM, with an average improvement of 24.4% (Figure 6, Table S8). Overall, such improvement was correlated with the improvement in ClinVar-gnomAD prediction AUROCs (Spearman correlation=0.76), and each score had at least 2.3% improvement for one of the predictive tasks. We note that for both prediction tasks, the improvement is more substantial for missense scores with low initial performance and less substantial for scores with high performance. For predicting ClinVar vs. gnomAD variants, scores in the bottom 50% for their initial performance had an average improvement of 25.4% compared to an average improvement of 6.3% for those in the top 50%. While for predicting DMS data, scores in the bottom 50% for their initial performance had an average improvement of 41.0% compared to an average improvement of 7.3% for those in the top 50%.

### Analyzing variant effect predictions from ProteinGym with MissenseHMM states

We next investigated the application of MissenseHMM to gain insights into predictions from 94 variant effect predictors that have predictions available in the ProteinGym database^39^. Many of these are recently developed predictors based on protein language models. We note that of these variant effect predictors, only ESM1b was included as input to MissenseHMM as the rest either did not have public predictions for all missense variants available or at least not in the dbNSFP database, which was the source of predictors for MissenseHMM. We specifically utilized the predictions for the zero-shot DMS substitutions benchmark from the ProteinGym database, which is based on predictions without observing the DMS labels in training and includes 58,335 variants from 96 DMS measurements coming from 81 proteins. We annotated these variants from the DMS experiments with MissenseHMM states. We then computed, for variants annotated to each state, the average ranks for each DMS measurement. We also computed for each state the average rank of each of the predictor scores across all variants in the state (Methods).

We observed that variants in different MissenseHMM states differ in their average ranks for DMS measurements, with states 1 and 2 showing the highest average ranks, corresponding to the highest pathogenicity measured by DMS (Figure 7, Table S9). In addition, the performance of the predictors on the DMS benchmark is associated with the specific states they prioritize. For example, predictors that assigned high scores to variants to the three states in Group 1 (states 1, 2 and 3) tended to perform better with the Spearman correlations across all scores’ leaderboard positions and their average ranks for these states ranged from 0.29 to 0.30 individually. In contrast, predictors that assigned high scores to variants in Group 2 states tended to have lower performance (average Spearman correlation -0.034 to -0.32), and the performance was lowest for the states in Group 3 (average Spearman correlation -0.35 to -0.53) and Group 4 (Spearman correlation of -0.47 and -0.49, Figure 7, Table S9).

**Figure 7:**
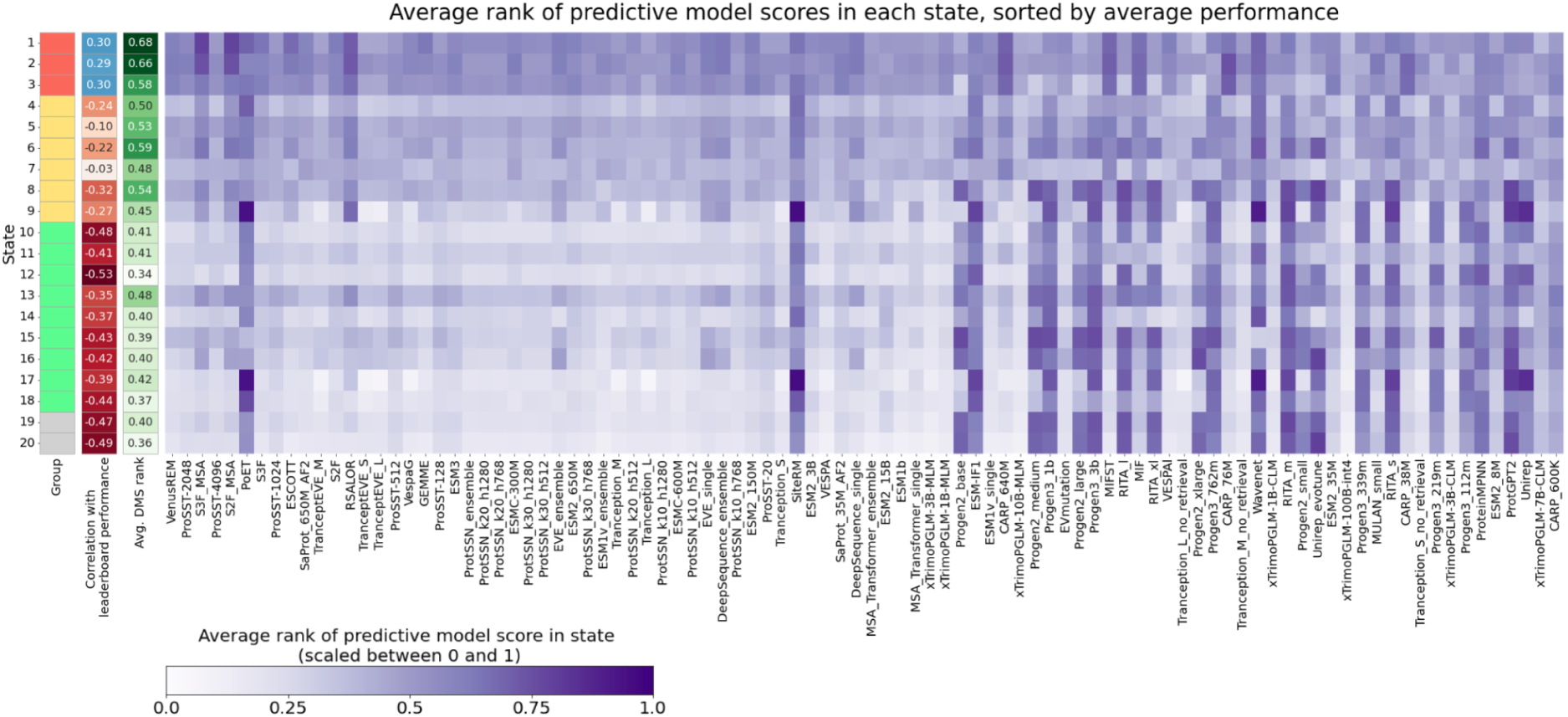
Average rank of predictor scores in each state, sorted by average performance across ProteinGym DMS experiments. Ranks are scaled between 0 and 1, with 0 representing the lowest rank and 1 representing the highest rank. Predictors have descending performance (i.e., leaderboard positions) from left to right. The second column from the left shows, for each state, the Spearman correlation between the average ranks of the predictors and the leaderboard positions of the predictors. Positive correlations indicate predictors with higher ranks in the state are more likely to place higher on the leaderboard, and vice versa. The third column from the left shows the average rank of DMS measurements in each state. The individual values corresponding to heatmap data is shown in Table S9.

Individual high-performing predictors also exhibit average variant score ranks across states that are highly correlated with those of the DMS measurements. Specifically, the 64 predictors whose average score rank distributions across the states had a Spearman correlation greater than 0.80 with those of the DMS measurements achieved a median leaderboard position of 36 out of 94, whereas the 30 predictors with correlations <0.80 had a median position of 70 out of 94 (two-sided Mann-Whitney U p-value=5.0x10^-6^, Figure S3). This indicates that strong performance in the evaluation is associated with a predictor prioritizing variants in the same states emphasized by the DMS. A notable exception is the PoET^40^ predictor, whose score ranks show only a weak correlation (0.094) with DMS measurements across the states, yet it achieves the eighth-best performance on the leaderboard.

## Discussion

Here we present MissenseHMM, an HMM-based approach to annotate missense variants through joint modeling of a wide array of missense scores. Unlike ensemble methods that integrate various missense scores into a single univariate output, MissenseHMM generates discrete states that can represent the combinatorial patterns of the scores that prioritize subsets of variants. We applied MissenseHMM to 43 missense scores in the dbNSFP database and analyzed a 20-state model where each state showed distinct emission patterns. We observed states showing prioritized variants across a large number of scores, states characterized by subsets of scores, states associated with variants prioritized by a single score, and states associated with variants generally not prioritized by any score. The MissenseHMM states exhibited distinct enrichments for external genomic annotations such as gene features, ChromHMM states, ConsHMM states. They also showed different compositions of amino acid substitutions. We also evaluated whether MissenseHMM annotations contain complementary information to individual scores. Our analysis demonstrated that, for a given individual score, the probability of a high-scoring variant being pathogenic associates with its MissenseHMM state.

Also, integrating MissenseHMM states with individual scores improved performance on pathogenicity prediction tasks, such as distinguishing ClinVar pathogenic variants from gnomAD benign variants and predicting the impact on protein function in experimental DMS data.

Additionally, we showed how MissenseHMM states provided insights into the performances of recently developed protein language model-based predictors on the ProteinGym benchmark.

We note MissenseHMM does have potential limitations and that it is designed to complement rather than replace traditional missense scores. One potential limitation of MissenseHMM is that the learned states do not fully retain the original information in the various scores used, some of which could be useful for variant prioritization. This is because MissenseHMM encodes the quantitative scores into binary input features using a quantile-based threshold and produces as output a discrete annotation of the genome. Another potential limitation of MissenseHMM annotations is that while they combine information from multiple scores, they do not preserve the flexibility of manual rule-based variant filtering that can combine multiple scores and which has been used previously^41^. Another limitation of MissenseHMM is that they may miss combinatorial relationships of interest or have less interpretable state distinctions, particularly for models with smaller and larger numbers of states, respectively. Overall, we expect scores from variant predictors to be preferred when the goal is fine-grained quantifications of pathogenicities of variants, while MissenseHMM could offer advantages when the goal is to obtain broad group-based categorizations of variants.

There are several potential future directions based on the MissenseHMM framework. One potential direction would be to investigate strategies to use MissenseHMM either alone or in conjunction with individual pathogenicity scores to analyze missense variants identified from large-scale genome sequencing studies. Another potential area of future investigation includes expanding the MissenseHMM model to incorporate more recent predictors, such as those evaluated by ProteinGym as genomewide predictions become available or more accessible for more of these predictors. Such updated models could further the understanding of the relationship between MissenseHMM state annotations and the performances of the various predictors on the ProteinGym benchmark. An additional direction for future work would be to adapt the MissenseHMM framework to the non-coding genome and integrate scores that prioritize variants genomewide^8,18,19,32^.

Overall, MissenseHMM provides a complementary view of many existing pathogenicity scores. We expect MissenseHMM and its annotations to facilitate the interpretation of missense variants through its integration of prioritization scores.

## Methods

### Input data and preprocessing

We obtained pathogenicity score predictions from 46 predictors for human nonsynonymous variants from the dbNSFP database v5.0a^6,7^, based on the human hg38 assembly. We subsetted to positions on the autosomes and chromosome X where only missense variants are present, leaving 70,842,617 missense variants from 25,665,962 unique positions. We used the ranked versions of scores provided by the database where each rank value is rescaled to the range [0, 1] where 0 represents the predicted least damaging effect and 1 represents the predicted most damaging effect. We removed the MutationTaster^42^ score due to its high missing rate (73%). We also removed phastCons470way_mammalian and phastCons100way_vertebrate scores^32^ because 62% and 55% of the variants, respectively, were tied for the highest score, leaving us scores from 43 predictors. Each variant was then represented as a vector *S = [S_1_, S_2_, S_3_…S_43_]* where *S_n_* = 1 if the variant had a value that was in the top *Q*% for score *n* across all missense variants and *S_n_* = 0 if it had a value was in the bottom (1 - *Q*)%. If more than *Q*% of the variants were tied for the highest score then they were all encoded as 1s. Following the default encoding method of the ChromHMM^26^ software, *S_n_* = 2 if the value for the missense variant was missing for score *n*. In our application of MissenseHMM here we set *Q* = 10.

### Model learning

To learn a MissenseHMM model, we first generated training sequences for the HMM. There was one sequence for each of the 19,563 protein-coding genes on the autosomes or chrX from the GENCODE database v46^41^. These sequences consisted of only positions with a missense variant. We note that each genomic position may have multiple missense variants with different alternative alleles. In our training sequences, we randomly selected one of the missense variants for each genomic position. We then created one binarized data file for each of the protein-coding genes, with the rows being the binary vectors of the selected variants ordered by their positions.

We next trained the HMM using the “LearnModel” command of the ChromHMM^26^ software using the gene-based binarized data files as input. We used the following flags: “-b 5000 -n 2000 -pseudo -d -1 -lowmem -p 4”. The “-b 1” flag specifies that the model’s resolution is 1bp. The “-n 2000” flag lets ChromHMM sample 2000 genes (about 10% of all genes) for each training iteration, reducing runtime, and the “-pseudo” flag adds pseudo-counts of 1 during parameter estimation to reduce sparsity caused by this subsampling. The “-d -1” flag allows ChromHMM to continue training even if the estimated likelihood decreases due to each training iteration using different subsamples of genes. The “-lowmem” flag reduces ChromHMM’s memory usage at the cost of some additional runtime. The “-p 4” flag specifies 4 parallel processors.

We trained models with 10 to 50 states in increments of 10. To assess how increasing the number of states affects the distinctiveness of emission patterns for individual states, we used the “CompareModels” command from ChromHMM^26^ to compute. Specifically, we used it to compute for each state in the 50-state model, the maximum correlation between that state and its best-matching counterpart in each smaller model. On average, each state in the 50-state model had a 0.83 maximum correlation with its best-matching state in the 20-state model, with 22 out of the 50 states having greater than 0.90 correlation and only 2 states having less than 0.20 correlation in comparison the corresponding values for the 10-state model were 0.72, 10, and 7 respectively reflecting individual states in the 50-state model that were not well captured with only 10 states (Figure S1, Table S10). While continuing to increase the number of states showed some quantitative improvements in the agreement with the 50-state model, the improvements were diminishing. Furthermore, as the 20-state model appeared to effectively capture the most meaningful variation in emission patterns and in the interest of increasing the interpretability of individual states, we chose the 20-state model for all subsequent analyses.

### Generating MissenseHMM state annotations for missense variants

After training, we used MissenseHMM’s resulting model to produce a state assignment for every missense variant based on estimating the most frequent state assignment for the missense variant over all possible sequences of missense variants for a gene. Finding the most common state of a missense variant over all possible sequences of a gene with *m* nucleotide positions directly would lead to up to 3*^m^* distinct sequences of missense variants. Instead to approximate the state assignments from such a procedure and avoid iterating through all these sequences, we generated “local combinations” in a similar manner as in a previous study^28^, that is for a target variant *v* at position *p_i_* we only considered *k* flanking positions upstream and downstream *(p_i-k_,…p_i-1_, p_i_, p_i+1_,…p_i+k_)* and generated *N* combinations where in each combination one missense variant was randomly selected for each of these positions except *p_i_*. We then applied the forward-backward algorithm to each of the *N* local combinations, obtaining for each the most probable state assignment for variant *v*. A mode of these *N* assignments was then used as the final assignment for variant *v*, with ties broken randomly. We generated state predictions for the 20-state model, with *k*=3 and *N*=9.

### Comparison to a model with uniform transition probabilities

To investigate the effect of the transition probabilities on model assignments, we created another 20-state MissenseHMM model where the emission probabilities are exactly the same as the original 20-state model but the transition probabilities were set to uniform (i.e. all transition probabilities were set to be 1/20=0.05). We applied this model to annotate the same set of missense variants as we did for the original model. Then, for each state in the original model, we computed the portion of variants assigned to each of the states in the model with uniform transition parameters.

### Compilation of external annotations

We obtained the following external annotations for conducting enrichment analysis:

- CpG islands annotations originally obtained from the UCSC genome browser^44^, were provided by default by ChromHMM v1.24^26^.
- Transcription start sites (TSS), transcription end sites (TES), and 2kb windows surrounding TSS (TSS2kb) in hg38 were obtained from the Gencode database, version v46^43^.
- 100-state universal ChromHMM state annotations in hg38 were obtained from Vu and Ernst^36^.
- 100-state ConsHMM state annotations of the hg38 reference sequence based on the 100-way vertebrate alignment were obtained from Arneson and Ernst^28,29^.

### Enrichments for external annotations

The fold enrichments were computed as *(B_s,a_ / B_s_) / (B_a_ / B_total_)* where *B_s,a_* was the number of variants in state *s* that also overlap with annotation *a*, *B_s_* was the number of variants in state *s*, *B_a_* was the number of variants overlapping annotation *a*, and *B_total_* was the total number of variants.

### Evaluating of predictive information in integration of MissenseHMM annotations with missense scores of ClinVar variants

To evaluate the additional predictive information in MissenseHMM annotations relative to individual missense scores’ in distinguishing pathogenic from putative benign variants, we curated benchmarking datasets using variants from ClinVar release 20240909^1^ and gnomAD v4.1^2^, labelled by dbNSFP. We selected two groups of variants:

1. ClinVar pathogenic: ClinVar variants labeled as either “pathogenic” or “likely_pathogenic”, with at least 1 gold star in review status, n=45,661
2. gnomAD singleton: gnomAD variants with minor allele count (MAC) = 1, n=6,017,743

We evaluated all 43 missense scores on the classification task of distinguishing between ClinVar pathogenic variants and gnomAD singletons, the latter we filtered to a subset that did not overlap with ClinVar pathogenic variants and then down-sampled to n=45,661. For each missense score we trained two separate logistic regression models for this task: the first model uses the missense score as its only feature, and the second model uses the missense score and the one-hot encoded MissenseHMM state assignments as its features. We evaluated the performance using 5-fold cross-validation and computed the overall AUROC scores.

### Evaluation of predictive information in integration of MissenseHMM annotations with missense scores of DMS datasets

To evaluate the additional predictive information in MissenseHMM annotations relative to individual missense scores’ for experimentally measured variant effects, we obtained DMS datasets from 36 proteins compiled by Livesey and Marsh^38^. Each dataset on average consisted of 2,429 variants, with the number of variants ranging from 138 variants to 10,015 variants.

Some of the datasets contained multiple DMS measurements. This resulted in a total of 96 measurements. We removed 10 proteins that were also used by the VARITY score for training, which left us with 26 proteins and 64 measurements. For each dataset, we partitioned the variants into five folds and used a cross-validation approach to evaluate each missense score’s predictive performance for each measurement. Specifically, for each combination of measurement, missense score and left-out fold, we trained two linear regression models on the remaining four folds. The first model predicts the DMS measurement of variants using one of the missense scores as its sole feature. The second model predicts the DMS measurement of variants using the missense score and the one-hot encoded MissenseHMM state assignment as its features. We then used both models to separately generate predictions for the variants in the left-out fold.

### Investigating the relationship between MissenseHMM states and predictors benchmarked by ProteinGym

To investigate the relationship between MissenseHMM states and the performance of protein mutation effect predictors benchmarked by ProteinGym, we downloaded files for 58,335 variants in 96 DMS experiments from 81 human proteins from the ProteinGym v1.3 zero-shot DMS substitutions benchmark^39^ (retrieved Apr. 27 2025) and analyzed them in the context of the 20-state MissenseHMM model. In the downloadable files, each variant had a DMS measurement and scores from 94 different predictors. To ensure comparability, both the DMS measurement and the scores from predictors were converted to ranks and scaled between 0 and 1. For the DMS measurements, the ranks were computed per-protein, with 0 and 1 representing the highest (least pathogenic) and lowest (most pathogenic) DMS measurements in the protein, respectively. For each of the predictor scores, the ranks were computed across proteins, also with 0 and 1 representing the highest (least pathogenic) and lowest (most pathogenic) scores. We first computed the leaderboard positions of each predictor by computing the average Spearman correlation between the predictor score ranks and the DMS measurement ranks, using the same procedure employed by ProteinGym except restricted to the subset of proteins that we considered. Then, for each state, we computed the Spearman correlation between the average rank of each predictor score in the state and the predictor’s position on the leaderboard in ascending order. For each predictor score, we computed the Spearman correlation between its average rank in individual states and the average rank of DMS in individual states.

## Declarations

### Ethics approval and consent to participate

Not applicable.

### Consent for publication

Not applicable.

### Competing interests

The authors declare that they have no competing interests.

### Availability of data and materials

The MissenseHMM software is available at https://github.com/ernstlab/MissenseHMM. The dbNSFP database v5.0a^6,7^ is available at https://www.dbnsfp.org/download. The ChromHMM^26^ software is available at https://ernstlab.github.io/ChromHMM/. 100-state universal ChromHMM state annotations in hg38 are from Vu and Ernst^36^ and are available at https://github.com/ernstlab/full_stack_ChromHMM_annotations. 100-state ConsHMM state annotations in hg38 are from Arneson and Ernst^28,29^ and are available at https://ernstlab.github.io/ConsHMMAtlas/. The DMS experimental data are from Livesey and Marsh^38^ and are available at https://doi.org/10.6084/m9.figshare.28295198. The ProteinGym benchmark data are from Notin *et al.*^39^ and are available at https://proteingym.org/benchmarks.

## Acknowledgements

We thank members of the Ernst lab for discussions and feedback. We thank Siddharth Naidu for preliminary work and discussions. We thank Tristan Bepler, Seon-Kyeong Jang, Zhengtong Liu, Sriram Sankararaman, and Noah Zaitlen for discussions.

## Funding

We acknowledge funding from US National Institutes of Health grants DP1DA044371, U01MH105578, R01MH109912, U01HG012079, and U01MH130995 (J.E.) and from a UCLA Dissertation Year Award (R.L.).

## Author contributions

R.L and J.E conceived & developed the methods, performed analyses and wrote the manuscript.

## Notes

https://github.com/ernstlab/MissenseHMM/

### Competing Interest Statement

The authors have declared no competing interest.

## References

1. Landrum, M. J. et al. ClinVar: public archive of relationships among sequence variation and human phenotype. Nucleic Acids Res. 42, D980–D985 (2014).

2. Karczewski, K. J. et al. The mutational constraint spectrum quantified from variation in 141,456 humans. Nature 581, 434–443 (2020).

3. Taliun, D. et al. Sequencing of 53,831 diverse genomes from the NHLBI TOPMed Program. Nature 590, 290–299 (2021).

4. Chen, S. et al. A genomic mutational constraint map using variation in 76,156 human genomes. Nature 625, 92–100 (2024).

5. Gao, H. et al. The landscape of tolerated genetic variation in humans and primates. Science 380, eabn8153 (2023).

6. Liu, X., Jian, X. & Boerwinkle, E. dbNSFP: A lightweight database of human nonsynonymous SNPs and their functional predictions. Hum. Mutat. 32, 894–899 (2011).

7. Liu, X., Li, C., Mou, C., Dong, Y. & Tu, Y. dbNSFP v4: a comprehensive database of transcript-specific functional predictions and annotations for human nonsynonymous and splice-site SNVs. Genome Med. 12, 103 (2020).

8. Rentzsch, P., Witten, D., Cooper, G. M., Shendure, J. & Kircher, M. CADD: predicting the deleteriousness of variants throughout the human genome. Nucleic Acids Res. 47, D886–D894 (2019).

9. Kim, S., Jhong, J.-H., Lee, J. & Koo, J.-Y. Meta-analytic support vector machine for integrating multiple omics data. BioData Min. 10, 2 (2017).

10. Jagadeesh, K. A. et al. M-CAP eliminates a majority of variants of uncertain significance in clinical exomes at high sensitivity. Nat. Genet. 48, 1581–1586 (2016).

11. Qi, H. et al. MVP predicts the pathogenicity of missense variants by deep learning. Nat. Commun. 12, 510 (2021).

12. Samocha, K. E. et al. Regional missense constraint improves variant deleteriousness prediction. Preprint at 10.1101/148353 (2017).

13. Sundaram, L. et al. Predicting the clinical impact of human mutation with deep neural networks. Nat. Genet. 50, 1161–1170 (2018).

14. Brandes, N., Goldman, G., Wang, C. H., Ye, C. J. & Ntranos, V. Genome-wide prediction of disease variant effects with a deep protein language model. Nat. Genet. 55, 1512–1522 (2023).

15. Cheng, J. et al. Accurate proteome-wide missense variant effect prediction with AlphaMissense. Science 381, eadg7492 (2023).

16. Jiang, T. T., Fang, L. & Wang, K. Deciphering “the language of nature”: A transformer-based language model for deleterious mutations in proteins. The Innovation 4, 100487 (2023).

17. Frazer, J. et al. Disease variant prediction with deep generative models of evolutionary data. Nature 599, 91–95 (2021).

18. Quang, D., Chen, Y. & Xie, X. DANN: a deep learning approach for annotating the pathogenicity of genetic variants. Bioinformatics 31, 761–763 (2015).

19. Ionita-Laza, I., McCallum, K., Xu, B. & Buxbaum, J. D. A spectral approach integrating functional genomic annotations for coding and noncoding variants. Nat. Genet. 48, 214–220 (2016).

20. Ioannidis, N. M. et al. REVEL: An Ensemble Method for Predicting the Pathogenicity of Rare Missense Variants. Am. J. Hum. Genet. 99, 877–885 (2016).

21. Li, B. et al. Automated inference of molecular mechanisms of disease from amino acid substitutions. Bioinformatics 25, 2744–2750 (2009).

22. Feng, B.-J. PERCH: A Unified Framework for Disease Gene Prioritization: HUMAN MUTATION. Hum. Mutat. 38, 243–251 (2017).

23. Alirezaie, N., Kernohan, K. D., Hartley, T., Majewski, J. & Hocking, T. D. ClinPred: Prediction Tool to Identify Disease-Relevant Nonsynonymous Single-Nucleotide Variants. Am. J. Hum. Genet. 103, 474–483 (2018).

24. Li, C., Zhi, D., Wang, K. & Liu, X. MetaRNN: differentiating rare pathogenic and rare benign missense SNVs and InDels using deep learning. Genome Med. 14, 115 (2022).

25. Dong, C. et al. Comparison and integration of deleteriousness prediction methods for nonsynonymous SNVs in whole exome sequencing studies. Hum. Mol. Genet. 24, 2125–2137 (2015).

26. Ernst, J. & Kellis, M. ChromHMM: automating chromatin-state discovery and characterization. Nat. Methods 9, 215–216 (2012).

27. Libbrecht, M. W., Chan, R. C. W. & Hoffman, M. M. Segmentation and genome annotation algorithms for identifying chromatin state and other genomic patterns. PLOS Comput. Biol. 17, e1009423 (2021).

28. Arneson, A., Felsheim, B., Chien, J. & Ernst, J. ConsHMM Atlas: conservation state annotations for major genomes and human genetic variation. NAR Genomics Bioinforma. 2, lqaa104 (2020).

29. Arneson, A. & Ernst, J. Systematic discovery of conservation states for single-nucleotide annotation of the human genome. *Commun*. Biol. 2, 248 (2019).

30. Wu, Y. et al. Improved pathogenicity prediction for rare human missense variants. Am. J. Hum. Genet. 108, 1891–1906 (2021).

31. Raimondi, D. et al. DEOGEN2: prediction and interactive visualization of single amino acid variant deleteriousness in human proteins. Nucleic Acids Res. 45, W201–W206 (2017).

32. Siepel, A. et al. Evolutionarily conserved elements in vertebrate, insect, worm, and yeast genomes. Genome Res. 15, 1034–1050 (2005).

33. Malhis, N., Jacobson, M., Jones, S. J. M. & Gsponer, J. LIST-S2: taxonomy based sorting of deleterious missense mutations across species. Nucleic Acids Res. 48, W154–W161 (2020).

34. Zhang, H., Xu, M. S., Fan, X., Chung, W. K. & Shen, Y. Predicting functional effect of missense variants using graph attention neural networks. *Nat*. Mach. Intell. 4, 1017–1028 (2022).

35. Dereli, O. et al. PHACTboost: A Phylogeny-Aware Pathogenicity Predictor for Missense Mutations via Boosting. Mol. Biol. Evol. 41, msae136 (2024).

36. Vu, H. & Ernst, J. Universal annotation of the human genome through integration of over a thousand epigenomic datasets. Genome Biol. 23, 9 (2022).

37. Henikoff, S. & Henikoff, J. G. Amino acid substitution matrices from protein blocks. Proc. Natl. Acad. Sci. 89, 10915–10919 (1992).

38. Livesey, B. J. & Marsh, J. A. Variant effect predictor correlation with functional assays is reflective of clinical classification performance. Genome Biol. 26, 104 (2025).

39. Notin, P. et al. ProteinGym: Large-Scale Benchmarks for Protein Fitness Prediction and Design. NIPS 23 Proc. 37th Int. Conf. Neural Inf. Process. Syst. 64331–64379 (2023).

40. Truong Jr., T. F. & Bepler, T. PoET: a generative model of protein families as sequences-of-sequences. NIPS 23 Proc. 37th Int. Conf. Neural Inf. Process. Syst. 77379–77415 (2023).

41. Zhang, T. et al. Heterologous survey of 130 DNA transposons in human cells highlights their functional divergence and expands the genome engineering toolbox. Cell 187, 3741–3760.e30 (2024).

42. Schwarz, J. M., Cooper, D. N., Schuelke, M. & Seelow, D. MutationTaster2: mutation prediction for the deep-sequencing age. Nat. Methods 11, 361–362 (2014).

43. Harrow, J. et al. GENCODE: The reference human genome annotation for The ENCODE Project. Genome Res. 22, 1760–1774 (2012).

44. Karolchik, D. The UCSC Genome Browser Database. Nucleic Acids Res. 31, 51–54 (2003).

