## Supplementary information for "MissenseHMM: state-based annotations for missense variants through joint modeling of pathogenicity scores"

### Supplementary Figures

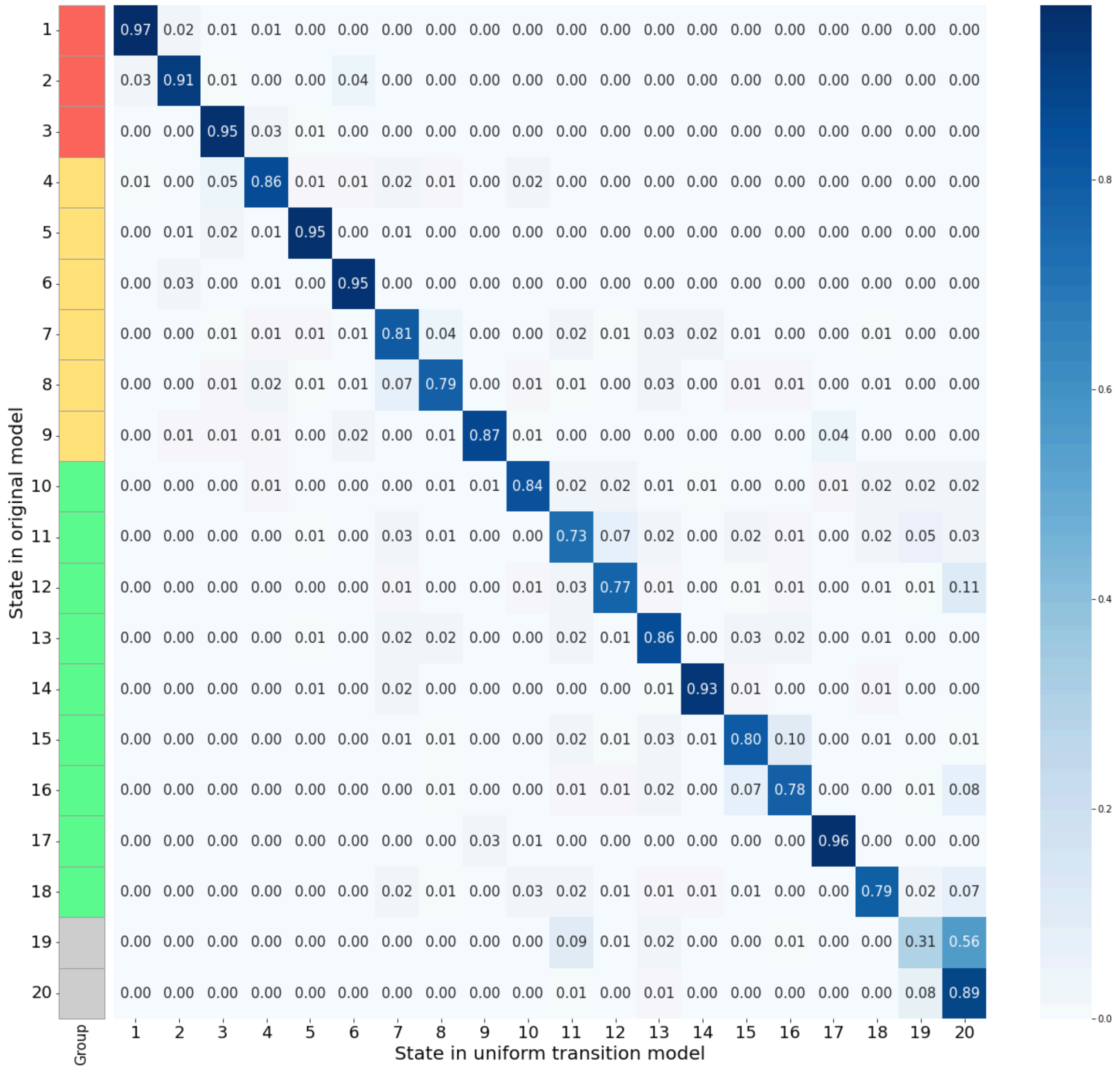


**Figure S1: Comparison between the original 20-state model and the uniform transition model.** For each state in the original model, the portion of variants assigned to each of the states in the uniform transition model are shown.

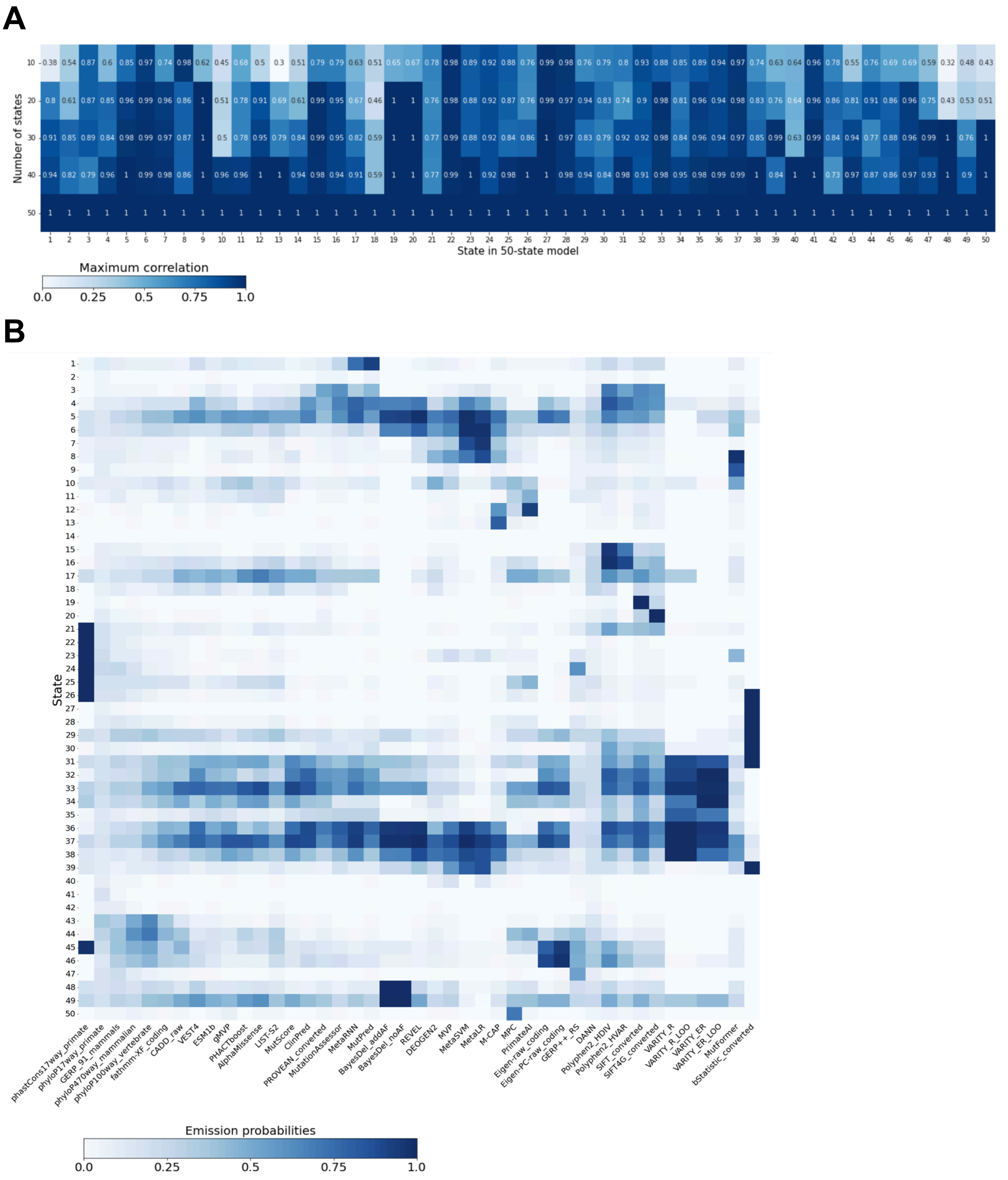


**Figure S2 CompareModels results using the 50-state model as reference. (A)** Each column represents a state in the 50-state model, and each row corresponds to a model with a different number of states. Heatmap cells indicate the maximum correlation between each state in the 50-state model and its best-matching counterpart in the smaller model. **(B)** Emission parameters for the 50-state model.

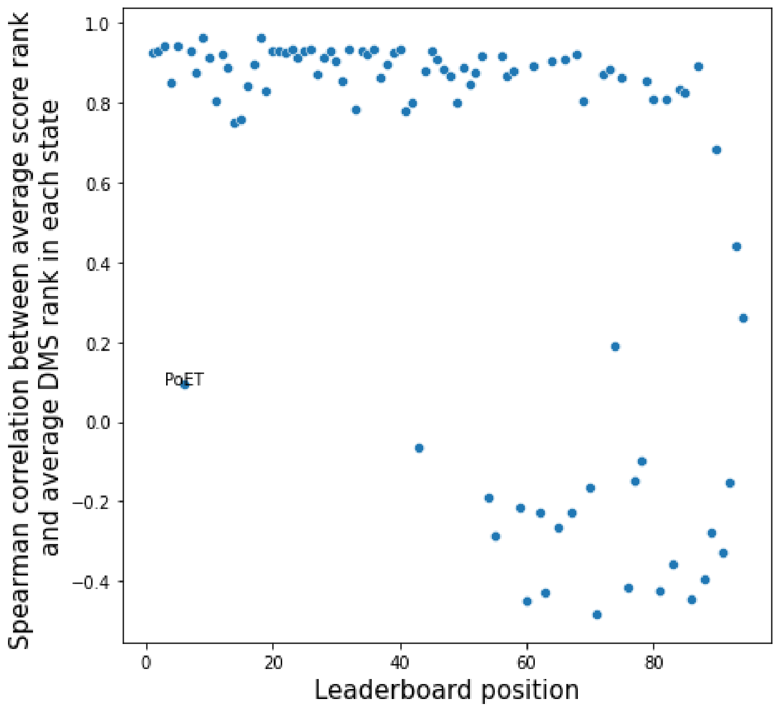


**Figure S3**: Across-state correlations between predictor scores and DMS measurements. X-axis shows the predictor's leaderboard positions, with 1 being the highest position. Y-axis shows the Spearman correlation between the predictor's average score rank in each state and the average DMS rank in each state (Methods).
